# Inhibition of the proton-activated chloride channel PAC by PIP_2_

**DOI:** 10.1101/2022.10.06.511171

**Authors:** Ljubica Mihaljević, Zheng Ruan, James Osei-Owusu, Wei Lü, Zhaozhu Qiu

**Author notes:** These authors contributed equally (Ljubica Mihaljević, Zheng Ruan).

## Abstract

Proton-Activated Chloride (PAC) channel is a ubiquitously expressed pH-sensing ion channel that regulates endosomal acidification and macropinosome shrinkage by releasing chloride from the organelle lumens. It is also found at the cell surface, where it is activated under pathological conditions related to acidosis and contributes to acid-induced cell death. However, the pharmacology of the PAC channel is poorly understood. Here, we report that phosphatidylinositol (4,5)-bisphosphate (PIP_2_) potently inhibits PAC channel activity. We solved the cryo-electron microscopy structure of PAC with PIP_2_ at pH 4.0 and identified its binding site, which, surprisingly, locates on the extracellular side of the transmembrane domain (TMD). While the overall conformation resembles the previously resolved PAC structure in the desensitized state, the TMD undergoes remodeling upon PIP_2_-binding. Structural and electrophysiological analyses suggest that PIP_2_ inhibits the PAC channel by stabilizing the channel in a desensitized-like conformation. To our knowledge, PAC is the first chloride channel reported to be inhibited by PIP_2_. Our findings identify PIP_2_ as a new pharmacological tool for the PAC channel and lay the foundation for future drug discovery targeting this channel.

## Introduction

Proton-activated chloride channel PAC is an evolutionarily conserved membrane protein with ubiquitous expression across different tissues. Since the recent discovery of its molecular identity (Yang et al., 2019, Ullrich et al., 2019), PAC has been implicated in important biological functions, such as endosomal trafficking and macropinocytosis (Osei-Owusu et al., 2021, Zeziulia et al., 2022). In the endosome, low luminal pH activates PAC to mediate chloride efflux from the lumen. Thereby, PAC actively regulates luminal acidification by depleting the counter ion, chloride, and preventing proton accumulation in the endosome (Osei-Owusu et al., 2021). During macropinocytosis, specialized endocytosis mediated by immune cells, PAC mediates the shrinkage of macropinosomes by releasing chloride into the cytoplasm (Zeziulia et al., 2022). In addition to localizing to the intracellular organelles, PAC also traffics to the plasma membrane, where it is involved several pathological conditions associated with acidosis. For example, upon ischemic stroke, PAC is activated by drops in tissue pH, allowing the entry of chloride into the cells. This subsequently causes cellular swelling and contributes to acid-induced brain injury (Osei-Owusu et al., 2020, Yang et al., 2019).

PAC is a homotrimer that forms a chloride-selective pore in the membrane, and it senses changes in pH via its large extracellular domain (ECD) (Ruan et al., 2020, Wang et al., 2022, Deng et al., 2021, Osei-Owusu et al., 2022a). PAC channel is closed at a neutral pH and becomes activated when the pH drops below 5.5 (Yang et al., 2019). The proton binding to the ECD is directly coupled with the channel opening in the transmembrane domain (TMD) (Osei-Owusu et al., 2022a). After prolonged exposure to pH 4.6 or below, the PAC channel slowly desensitizes (Osei-Owusu et al., 2022). The desensitization of PAC is pH-dependent, i.e., under more acidic conditions, desensitization is increased with faster kinetics (Osei-Owusu et al., 2022). This is regulated by several key residues localized at the ECD–TMD interface (Osei-Owusu et al., 2022). For example, the E94R mutant displays fast desensitization even at pH 5.0, when the wild-type channel does not exhibit obvious current decay (Osei-Owusu et al., 2022b). In addition to the resting and desensitized structures (Ruan et al., 2020), an open conformation of PAC was recently reported (Wang et al., 2022). During the transition between different channel conformations, major structural rearrangements of PAC happen inside the lipid bilayer (Ruan et al., 2020, Wang et al., 2022). The notion that the membrane shapes ion channel function and structure is now widely accepted. Lipid composition and the thickness of the membrane can directly control or fine-tune the gating of certain ion channels (Rosenhouse-Dantsker et al., 2012). However, whether the PAC channel is regulated by lipids is unknown.

The most common and best-studied lipid regulator of ion channel function is phosphatidylinositol (4,5)-bisphosphate (PIP_2_). PIP_2_ is a negatively charged phospholipid, predominantly found in the inner leaflet of the plasma membrane (Suh & Hille, 2008), with few reports that it can localize to the outer leaflet as well (Gulshan et al., 2016, Yoneda et al., 2020). Although it accounts for less than 1% of the total phospholipids in the plasma membrane, it is a principal signaling molecule and an essential cofactor for ion channel function (Hansen, 2015; Suh & Hille, 2008). PIP_2_ binds to ion channels directly and modulates their function by facilitating channel opening, preventing current rundown/desensitization, or inhibiting channel activity (Gada & Logothetis, 2022; Suh & Hille, 2008). At least ten different ion channel families are dependent on PIP_2_ for their activity, of which only one chloride channel family, TMEM16A, is known to require PIP_2_ for channel opening. Additionally, a handful of ion channels are reported to be inhibited by PIP_2_ (Gada & Logothetis, 2022; Suh & Hille, 2008).

Major breakthroughs have been made in recent years to characterize the structure and function of the PAC channel in biology and disease, but its regulation by endogenous molecules remains largely unexplored. To date, arachidonic acid and pregnenolone sulfate are the only reported potential biological inhibitors of the acid-induced chloride currents (*I_Cl,H_*) mediated by PAC, yet it is unknown if their mechanism is direct or indirect (Sato-Numata et al., 2016, Okada et al., 2021). Here, we show that the PAC channel is inhibited by the direct binding of PIP_2_ to the protein, representing the first known chloride channel inhibited by PIP_2_. Furthermore, we elucidate the molecular mechanism by which PIP_2_ inhibits PAC by stabilizing a desensitized-like conformation of the channel.

## Results

### PIP_2_ inhibits PAC channel activity

Considering the widespread influence of PIP_2_ on ion channel function, we hypothesized that PIP_2_ could potentially regulate the PAC channel. To test whether PIP_2_ modulates PAC activity, we applied a soluble version of PIP_2_ lipid, dioctanoyl phosphatidylinositol 4,5 bisphosphate (diC_8_-PIP_2_), to HEK293 cells, which endogenously express the acid-induced chloride currents. Whole-cell *I*_Cl,H_ were detected in real-time, by perfusing the cells with an acidic solution at pH 5.0, followed by an application of 10 μM diC_8_-PIP_2_. Immediately upon adding PIP_2_, there was a rapid drop in *I*_Cl,H_ followed by a steady decline (**Figure 1A**). The time constant, Tau, was calculated using a one-phase decay equation, and for PAC inhibition by PIP_2_ at pH 5.0 Tau was 56 ± 14.82 s, n = 7. Furthermore, this effect was reversible by washing out the soluble lipid from the cell membrane with a pH 5.0 solution (**Figure 1A**). ~37% of the initial current amplitude was detectable after PIP_2_ perfusion for 150 s (**Figure 1B, C**). This short timescale of PIP_2_ action on PAC activity indicates that its effect is most likely direct. Further supporting this, PIP_2_ inhibited *I*_Cl,H_ in a dose-dependent manner, with half-maximal inhibition, IC_50_, of 4.9 μM (**Figure 1D**). The Hill coefficient value was equal to 1.57, suggesting two or more potential binding sites on the protein.

**Figure 1.**
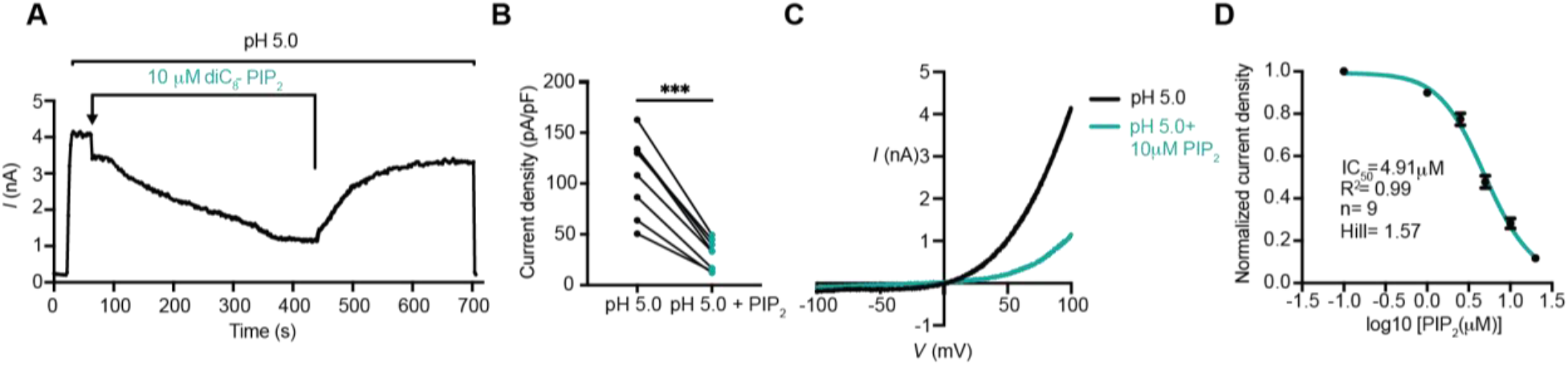
PIP_2_ inhibits the PAC channel activity. **(A)** Representative whole-cell current trace (2s/sweep) showing inhibition of endogenous PAC currents by bath perfusion of soluble diC_8_-PIP_2_ at 10 μM concentration. **(B)** PAC current densities before and after application of PIP_2_ for 150 s. Statistical significance was determined using a two-tailed Student’s paired *t* test. ***p < 0.0001. **(C)** Representative *I/V* curve of pH 5-induced PAC current before and after PIP_2_ treatment. **(D)** Dose-dependent inhibition of the PAC channel by PIP_2_ yielded a half-maximal inhibition, IC_50_, of 4.91 μM with a Hill slope of 1.57. Bars are reported as mean ± SEM.

At neutral pH, the PAC channel is in its resting/closed state, and it is activated/open when the pH drops below 5.5 at room temperature (Yang et. al, 2019). To examine if PIP_2_ exerts its effect on the PAC channel in its closed or open state, we pre-treated the cells with soluble PIP_2_ at pH 7.3, and then activated the channel with acid. *I*_Cl,H_ amplitude at pH 5.0 did not show any significant difference before and after perfusion of PIP_2_ at the neutral pH (**Figure S1A,B**). This result suggests that PIP_2_ may not act on the resting state of PAC and is only effective once the channel undergoes proton-induced activation or the subsequent desensitization.

### Phosphates and the acyl chain synergistically contribute to PIP_2_-mediated PAC inhibition

To test if there is a preference among different phosphatidylinositol lipids, we used soluble (diC_8_) versions of lipids at 10 μM concentration and compared their inhibitory effects on PAC. PI(3)P (phosphatidylinositol 3-phosphate) with a single phosphate on its inositol headgroup, inhibited PAC significantly less than either bisphosphonates, PI(4,5)P_2_ or PI(3,5)P_2_ (phosphatidylinositol (3,5)-bisphosphate) (**Figure 2A**). The additional phosphate on PIP_3_ (phosphatidylinositol (3,4,5)-trisphosphate) further lowered the IC_50_ to 3 μM (**Figure 2B**). Therefore, to reach potent inhibition, a minimum of two phosphates on the inositol headgroup are required. This is additionally supported by a modest inhibitory effect of PI (phosphatidylinositol) that does not have any active headgroup phosphates (**Figure 2C**). Interestingly, IP3 (inositol 1,4,5-trisphosphate), a triple-phosphorylated inositol headgroup without an acyl chain, displayed a similarly modest inhibition on PAC as PI (**Figure 2D**). Phosphates on the headgroup are therefore necessary, but not sufficient for PAC inhibition, indicating that the lipid chain contributes to inhibitory properties of PIP_2_ as well. diC_8_-diacyl-glycerol (DAG), the lipid chain without inositol head, had no inhibitory effect on PAC (**Figure 2D**). Acyl chain alone was therefore not sufficient to inhibit PAC.

**Figure 2.**
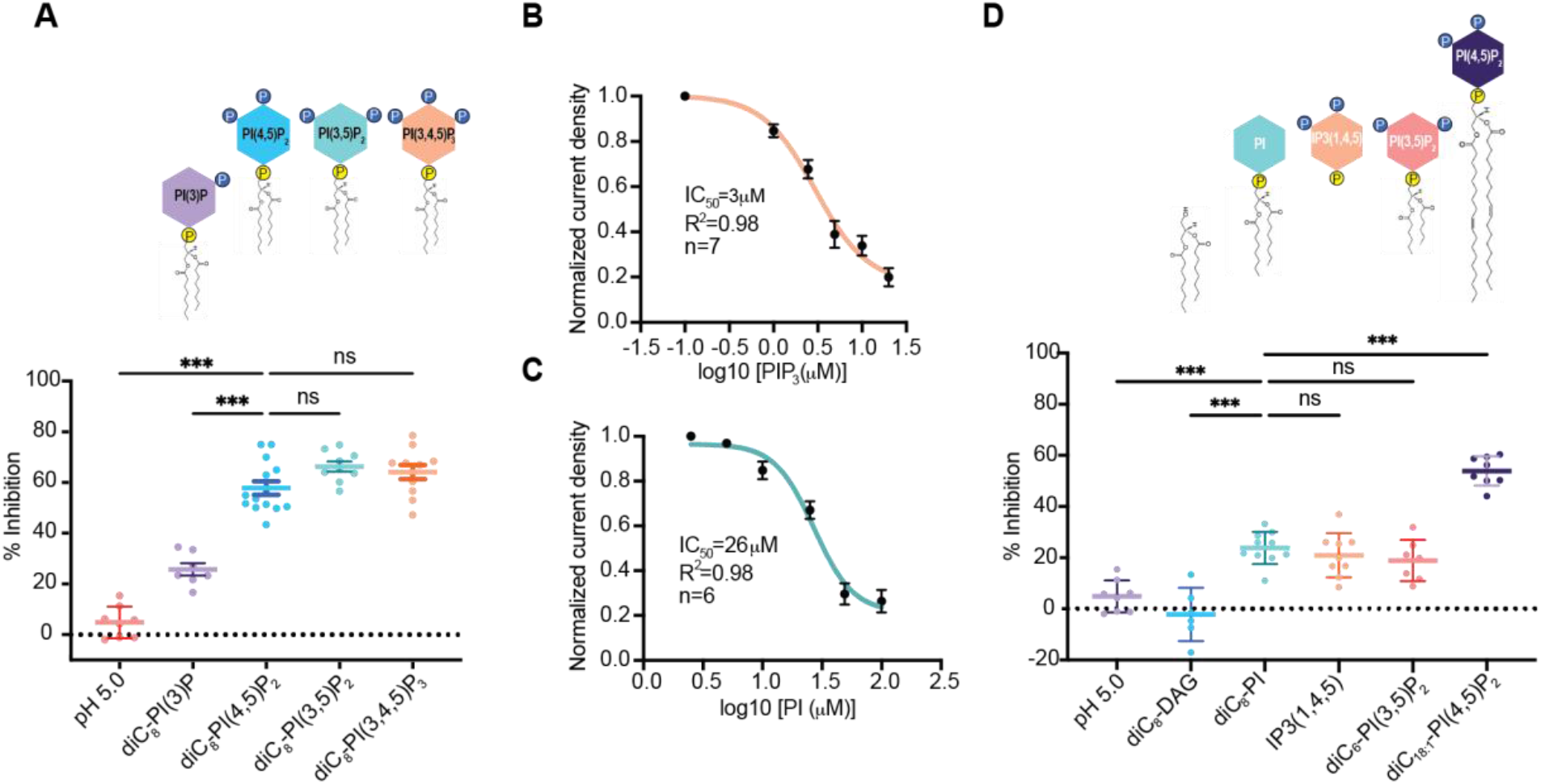
Phosphates and acyl chain length synergistically contribute to PAC inhibition by PIP_2_. **(A)** Percent inhibition of pH 5-induced PAC current by different diC_8_-phosphatidylinositol lipids. PI(3)P inhibited PAC significantly less than PI(4,5)P_2_, PI(3,5)P2 and PI(3,4,5)P3.There was no significant difference between PI(4,5)P_2_, PI(3,5)P2 and PI(3,4,5)P3. Statistical significance was determined using ordinary one-way ANOVA with Dunnett post hoc test. ***p < 0.0001; ns: not significant. Bars are reported as mean ± SEM. **(B)** Dose-dependent inhibition of PAC current by PIP_3_. PIP_3_ is the most potent inhibitor of PAC with half-maximal inhibition, IC_50_, of 3 μM. Bars are reported as mean ± SEM. **(C)** Dose-dependent inhibition of PAC current by phosphatidylinositol (PI). IC_50_ of PI is higher than (26 μM) than PIP_2_ (5 μM) or PIP_3_ (μM). Bars are reported as mean ± SEM. **(D)** Acyl chain length contributes to PAC inhibition by PIP_2_. Diacyl glycerol (DAG), which has just an acyl chain without an inositol headgroup didn’t inhibit PAC. diC_8_-PI, IP3(1,4,5) and diC_6_-PI(3,5)P_2_ displayed modest inhibition of PAC. Natural diC_18:1_-PI(4,5)P_2_ potently inhibited the PAC channel. All phosphatidylinositols were perfused at 10 μM concentration, except the less soluble diC_18:1_-PIP_2_ at 20 μM. Statistical significance was assessed using ordinary one-way ANOVA with Dunnett post hoc test. ***p < 0.0001; ns: not significant. Bars are reported as mean ± SEM.

Next, we examined the inhibitory effect of phosphatidylinositol lipids with varying chain lengths. PIP_2_ with two carbons less on its acyl chain, diC6-PIP_2_, was significantly less potent than diC_8_-PIP_2_ in inhibiting *I*_Cl,H_ (**Figure 2D**). On the other hand, PIP_2_ with a naturally occurring, long, lipid chain, diC_18:1_-PIP_2_, displayed a strong inhibition on the PAC channel (**Figure 2D**). Based on these results, we conclude that an acyl chain with a minimum of 8 carbons is required for potent inhibition of PAC by PIP_2_. Together, the number of phosphates on the inositol headgroup and lipid chain length synergistically contribute to the inhibitory potency of PIP_2_ to the PAC channel.

### Cryo-EM structure reveals the PIP_2_ binding site on the PAC channel

PIP_2_ often binds to motifs on the intracellular side of ion channels, which contain positively charged residues that directly interact with the negatively charged phosphates on its inositol head. Indeed, the C-terminus of TM2 has several positively charged residues, including K325, K329, K333, R335, K336, R337, K340, R341, R342, that we focused on initially and studied for their impact on PIP_2_ sensitivity. (**Figure S2A**). Mutating single or triple lysine and arginine residues to alanine or making a 10-residue deletion at the C-terminal domain did not affect PIP_2_-mediated PAC inhibition (**Figure S2B**). In addition, when diC_8_-PIP_2_ was applied to the cells through an intracellular solution in the patch pipette, at a physiological concentration of 10 μM, there was no detectable change in the PAC current amplitude (**Figure S2C**). Similarly, *I*_Cl,H_ remained intact when endogenous PIP_2_ was depleted from the inner membrane leaflet using 100 μg/ml Poly-l-Lysine (PLL) in the patch pipette (**Figure S2D**). These results are surprising because endogenous PIP_2_ is known to be almost exclusively localized to the inner leaflet of the plasma membrane. Thus, the effect we observed with the perfusion of exogenous PIP_2_ may occur via inhibition of the PAC channel through a potentially unconventional mechanism.

To reveal the mechanism underlying PIP_2_ inhibition, we solved the cryo-EM structure of PAC in nanodiscs with 0.5 mM diC_8_-PIP_2_ at pH 4.0 to an overall resolution of 2.71 Å (**Figure 3A, Table S1**). The structure adopts a conformation similar to the previously reported desensitized state at low pH (Ruan et al., 2020). However, a strong branched lipid density is observed on the unsharpened cryo-EM map of the PAC channel in the outer membrane leaflet, between TM1 and TM2 of adjacent subunits (**Figure 3A, Figure S3, S4**). The density appears to be a PIP_2_ molecule based on the shape of the cryo-EM map density, although it is by no means unambiguous due to the limited local map resolution and the seemingly high flexibility of the putative lipid molecule associated with this density. After fitting a PIP_2_ molecule into this density (**Figure 3B**), we found that the phosphatidyl group is reasonably well defined, with its phosphate group forming a salt bridge interaction with R93 and its two acyl tails interacting with a number of hydrophobic residues on both transmembrane helices, including F92 and W304 (**Figure 3C)**. In contrast, the inositol-4,5-bisphosphate moiety is not resolved and therefore not modeled, probably because it does not make close contact with nearby residues and is therefore flexible. Nevertheless, the local biochemical environment of the site is consistent with PIP_2_ binding, in which the negatively charged inositol-4,5-bisphosphate head group is surrounded by several positively charged residues, including K97, K106, and K294 (**Figure 3C)**. Overall, the putative PIP_2_ binding site is in accordance with our observation that the higher number of negatively charged phosphates, as well as the presence of an acyl chain, contribute to stronger channel inhibition by PIP_2_ (**Figure 2**).

**Figure 3.**
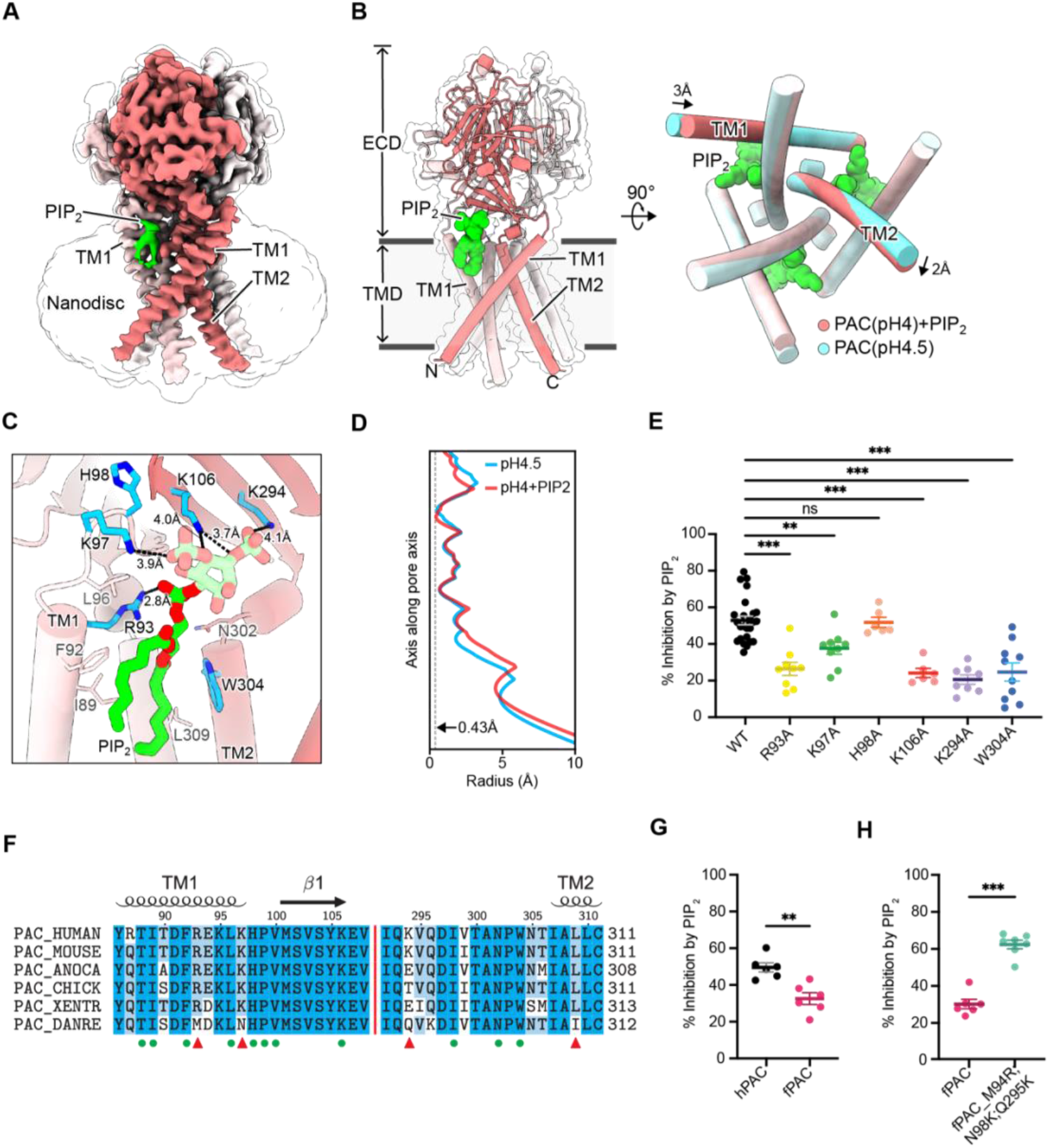
PIP_2_ binds directly to the PAC channel. **(A)** Cryo-EM structure of PAC channel at pH 4.0 with bound PIP_2_. One subunit is shown in red with TM1 and TM2 labelled. Density corresponding to PIP_2_ is colored in green. Inositol-4,5-bisphosphate group is not resolved in the Cryo-EM map and thus not modeled in the deposited structure; the model of the inositol-4,5-bisphosphate group in the figure is only to show its local biochemical environment. **(B)** The structural model of PAC channel at pH 4.0 with bound PIP_2_ in side view (left) and bottom-up view (right). For comparison, the PAC channel at pH 4.5 without PIP_2_ is shown in cyan for the bottom-up view (right). **(C)** A close-up view of the PIP_2_ binding site. Residues close to the putative PIP_2_ binding site are shown in stick representation. Residues tested in the study, including R93, K97, H98, K106, K294, and W304 are colored in blue. Distances between positively charged residues and the phosphate group are indicated. The inositol-4,5-bisphosphate group (shown in tinted color) is not resolved in the Cryo-EM map and thus not modeled in the deposited structure. The model of the inositol-4,5-bisphosphate group in the figure is only to show its local biochemical environment. **(D)** The pore profile of PAC at pH 4.0 with PIP_2_ and at pH 4.5 without PIP_2_. The smallest radius along the pore axis is 0.43 Å, suggesting that both structures are impermeable to chloride ions. **(E)** Mutating PIP_2_-binding residues to alanine significantly decreases % inhibition by diC_8_-PIP_2_. Statistical significance was assessed using one-way ANOVA with Dunnett post hoc test. ***p < 0.0001; **p < 0.01; ns: not significant. Bars are reported as mean ± SEM. **(F)** Multiple sequence alignment of several PAC orthologs. Key residues that form the PIP_2_ binding site are labelled using green dots. Three binding site residues that are not conserved in zebrafish PAC (PAC_DANRE) are indicated by red triangles. **(G)** Percent inhibition (mean ± SEM) of hPAC or fPAC current at pH 5 by 10 μM diC_8_-PIP_2_. Zebrafish PAC (fPAC) shows significantly less inhibition by PIP_2_ compared to human PAC. Statistical significance was determined using a two-tailed Student’s unpaired *t* test. **p < 0.01. **(H)** Mutating zebrafish PAC residues to the corresponding human PAC residues significantly increases the inhibition by PIP_2_ in comparison to the wild-type zebrafish PAC. Statistical significance was determined using a two-tailed Student’s unpaired *t* test. ***p < 0.0001. Bars are reported as mean ± SEM.

Small, but notable conformational changes are observed in the transmembrane helices (TM1 and TM2) upon PIP_2_ binding. Specifically, TM1 tilts inside by 3 Å which causes a concerted rotation motion of TM2 (**Figure 3B**). The pore radius profile is similar to the desensitized state of PAC without PIP_2_, with the smallest radius of 0.43 Å (**Figure 3D**). Therefore, the PIP_2_-bound conformation also represents a non-conductive state (**Figure 3A-C).** To validate our structural model, we carried out site-directed mutagenesis and patch-clamp electrophysiological experiments. Mutation of any residues R93, K97, K106, K294, and W304 to alanine significantly relieved the inhibition of PAC by PIP_2_, confirming that this was indeed its binding site on the channel (**Figure 3E**). We also examined an adjacent residue, H98, which is not at a distance from the binding site that would allow direct interaction with PIP_2_. As expected, H98A mutant was still sensitive to PIP_2_ inhibition (**Figure 3E**), suggesting that PIP_2_ specifically recognizes the binding pocket observed in our structure. None of the mutations we tested affected the PAC channel activity, as indicated by the normal current densities (**Figure S2E**).

Putative PAC PIP_2_-binding residues are conserved amongst higher vertebrates. On the contrary, in zebrafish *(Danio rerio),* several PIP_2_-binding site residues are different, including M94, N98, and Q295 (**Figure 3F**). Interestingly, the zebrafish PAC channel (fPAC) was significantly less inhibited by PIP_2_ compared to the human PAC (hPAC) (**Figure 3G**). To test if the reduced PIP_2_ sensitivity of zebrafish PAC is due to these amino acid differences, we used site-directed mutagenesis to convert zebrafish residues to the corresponding ones of the human PAC channel. Interestingly, zebrafish triple mutant M94R, N98K, Q295K (fPAC numbering) showed a significant increase in PIP_2_ inhibition when compared to the wild-type channel (**Figure 3H, Figure S2F**). This further substantiates our finding that the PIP_2_ binding site on the PAC channel is located in the outer membrane leaflet.

### PIP_2_-mediated PAC inhibition correlates with the degree of channel desensitization

Since PIP_2_-bound PAC structure resembles the desensitized state and the PAC channel desensitization is regulated by pH, we sought to examine how pH may influence PIP_2_-mediated PAC inhibition by applying diC_8_-PIP_2_ at a pH range from 4.0 to 5.2 (**Figure 4A-C**). The percentage of PAC inhibition by PIP_2_ increased with a decrease in pH (**Figure 4D**). Furthermore, the time to reach maximal inhibition was also pH-dependent, with significantly faster kinetics of PIP_2_-mediated PAC inhibition at the lower pHs (**Figure 4E**). These results suggest that PIP_2_ inhibition is more effective when the PAC channel is already poised towards the desensitized state under more acidic conditions.

**Figure 4.**
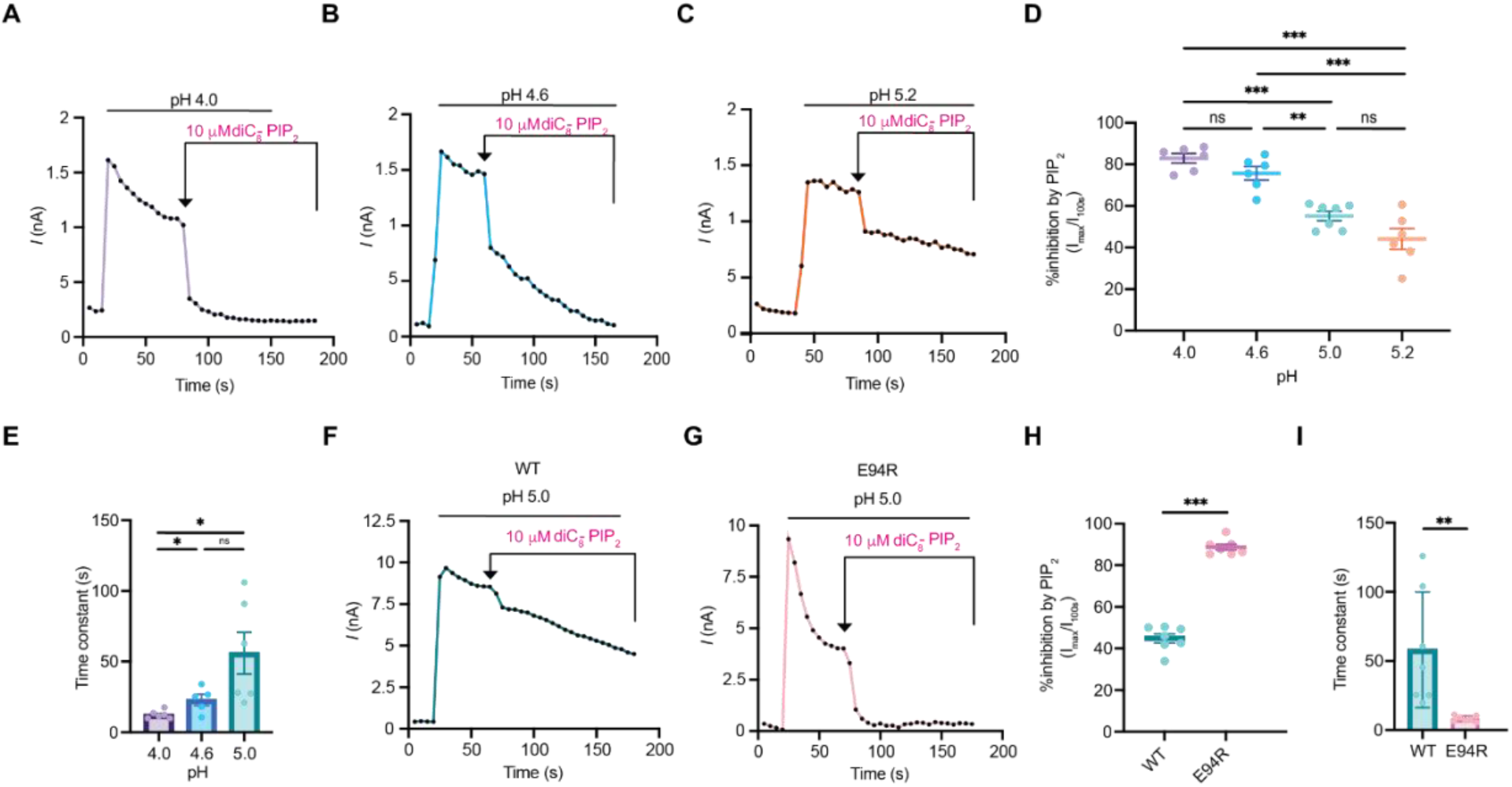
PIP_2_-mediated PAC inhibition correlates with the degree of channel desensitization. **(A), (B), (C)** Representative current traces (5s/sweep) of endogenous PAC currents at pH 4.0, 4.6 and 5.2 treated with 10 μM diC_8_-PIP_2_. Note that, PAC desensitizes more at pH 4.0 than at pH 4.6, while there is no obvious desensitization at pH 5.2. **(D)** The percent of PAC inhibition (mean ± SEM) by PIP_2_ decreases with an increase in pH. I_max_ is the current amplitude right before applying PIP_2_, after desensitization reaches a plateau. At 100 s after diC_8_-PIP_2_ perfusion, PAC inhibition by PIP_2_ reached 83% at pH 4.0, 76% at 4.6, 55% at 5.0, and 44% at the pH 5.2. Statistical significance was determined using one-way ANOVA with Dunnett post hoc test. ***p < 0.0001; **p < 0.01; ns: not significant. **(E)** The time constant, Tau, was calculated using a one-phase decay equation and plotted as mean ± SEM for pH 4.0, 4.6 and 5.0 (for pH 5.2, time constant fits were below R^2^=0.8, therefore considered an unreliable). The kinetics of PIP_2_ inhibition is slower at higher pH. Statistical significance was determined using a two-tailed Student’s unpaired *t* test. *p < 0.05; ns: not significant. **(F), (G)** Representative current traces (5s/sweep) of overexpressing PAC WT and E94R at pH 5.0 treated with 10 μM diC_8_-PIP_2_. **(H)** Percent inhibition (mean ± SEM) of PAC WT and E94R current at pH 5.0 100s after perfusion of 10 μM diC_8_-PIP_2_. Statistical significance was determined using one-way ANOVA with Dunnett post hoc test. ***p < 0.0001. **(I)** The time constant, Tau, was calculated using a one-phase decay equation and plotted as mean ± SEM for for PAC WT and E94R mutant. E94R displays faster kinetics of inhibition by PIP_2_ compared to WT at pH 5.0. Statistical significance was determined using a two-tailed Student’s unpaired *t* test. **p < 0.01.

We recently showed that reversing the charge of E94 residue to E94R induces PAC desensitization, even at pH 5.0 (**Figure 4F,G**) (Osei-Owusu et al., 2022b). Structurally, E94 is located in TM1, facing the opposite side of the PIP_2_ binding pocket. Therefore, E94 mutation is unlikely to affect PIP_2_ binding directly, representing an ideal candidate to test if there is a correlation between PIP_2_ inhibition and channel desensitization. Indeed, we found that PIP_2_ exerted a much higher degree of inhibition on the E94R mutant than the WT PAC channel (**Figure 4H**). In addition, the E94R mutant reached its maximal inhibition by PIP_2_ significantly faster than the WT channel (**Figure 4I**). Because E94 is distal to the PIP_2_ binding site, these effects are most likely due to the altered conformational dynamics toward desensitization. Together with its pH-dependency, our data suggest that PIP_2_ inhibition is more effective when the PAC channel is more prone to becoming desensitized.

## Discussion

PAC is a novel chloride channel and its pharmacology is still poorly studied. Here, we showed that PIP_2_ binds to and potently inhibits the PAC channel with an IC_50_ of ~4.9 μM. This value is comparable to the EC50 for PIP_2_ activation of TMEM16A—the only other chloride channel known to directly bind PIP_2_ prior to this work—and other PIP_2_-regulated cation channels, such as inwardly rectifying potassium channel K_ir_ (Le et al., 2019). Our structural analysis further showed that the PIP_2_ binding site in PAC is located on the extracellular side of the TMD, unlike other ion channels known to be regulated by PIP_2_, which bind PIP_2_ on the intracellular side of the TMD.

Our data indicate that the degree and kinetics of PIP_2_-mediated PAC inhibition depend on channel desensitization. The prevalence of desensitized PAC state at low pH (**Figure 4A**) and in the E94R mutant (**Figure 4G**) facilitates the inhibitory effect of PIP_2_ on PAC. PIP_2_-mediated PAC inhibition under these conditions is more effective likely because the desensitized conformation becomes more prevalent and accessible. Contrarily, under less acidic conditions (**Figure 4C,D)** most of the channels adopt the open/resting states, and the presence of PIP_2_ could only slowly shift the equilibrium towards the desensitized state. This is reflected in the larger inhibition time constants at less acidic pHs and in WT when compared to those at lower pHs and in the E94R mutant (**Figure 4E, I**). Together, our results suggest that PIP_2_ achieves its inhibitory effect by promoting the desensitized state of the PAC channel. In our proposed model (**Figure 5**), PIP_2_ selectively binds to the desensitized conformation of PAC and lowers the energy barrier (ΔG) for PAC to transition to the desensitized state. By rearranging the conformational landscape of PAC towards the desensitized state, PIP_2_ potently inhibits the PAC channel.

**Figure 5.**
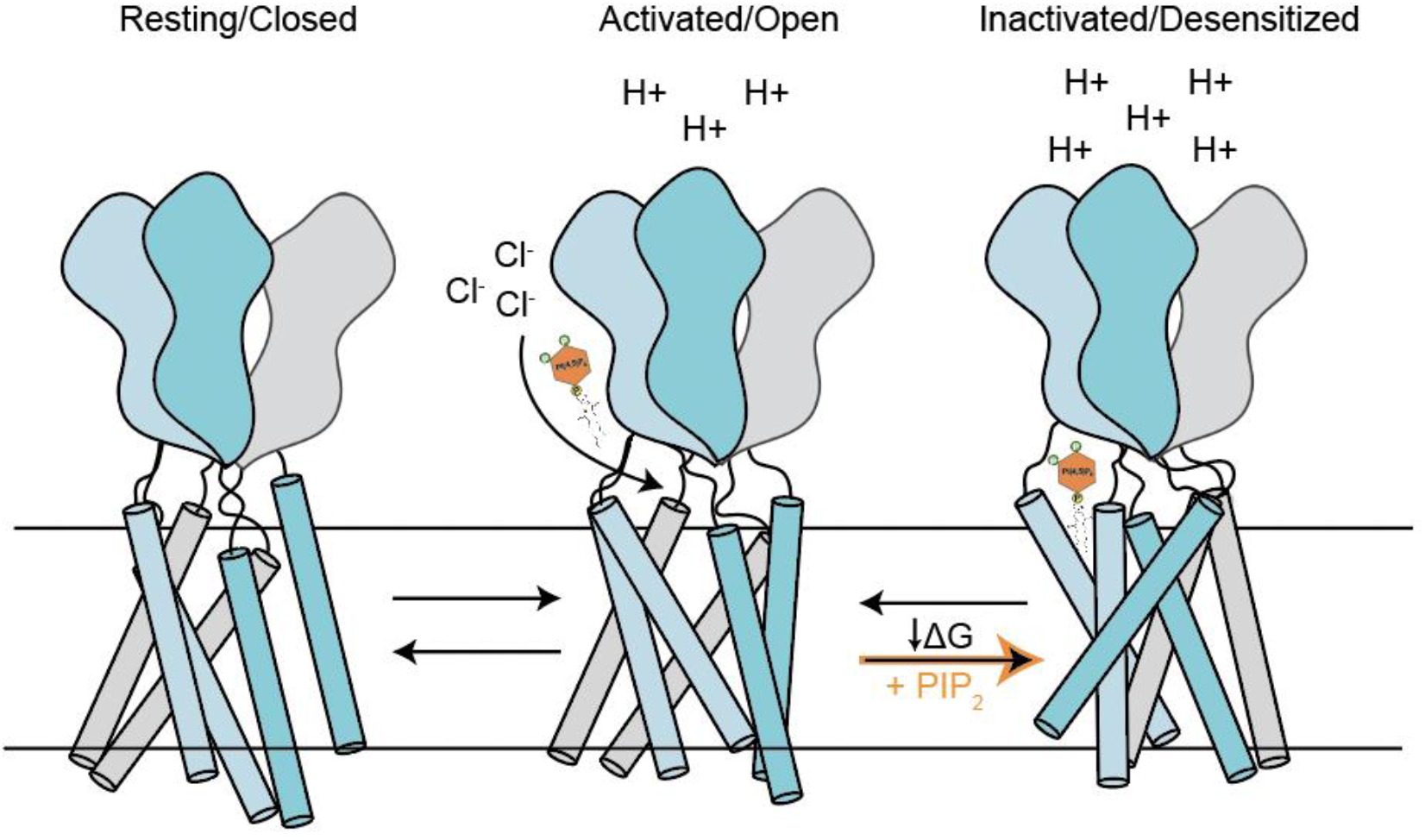
A proposed model of PAC inhibition by PIP_2_. Once PAC is activated, PIP_2_ enters the channel. The PIP_2_ binding shifts the conformational landscape of the channel, lowering the energy barrier (ΔG) towards the PAC desensitized state. Subsequently, PIP_2_ locks the PAC channel in its desensitized conformation, leading to its inhibition.

Furthermore, we characterized pharmacological properties that contribute to phosphatidylinositol inhibitory potency on PAC. PIP_3_ displayed the strongest inhibition of PAC, while IP3 had a negligible effect (**Figure 2**). A higher number of phosphates and a longer acyl chain increase the inhibitory potency of the lipid. This indicates that the negative charge on the inositol head, as well as acyl chain insertion into the membrane, synergistically contribute to the binding and stabilization of the desensitized channel state.

PIP_2_ is primarily localized in the cytosolic side of the plasma membrane. However, a few studies report that PIP_2_ can be detected on the extracellular side (Gulshan et al., 2016; Yoneda et al., 2020). In RAW264.7 macrophages and baby hamster kidney (BHK) fibroblasts, ATP-binding cassette transporter A1 (ABCA1) facilitates the redistribution of PIP_2_ from the inner to the outer leaflet of the plasma membrane (Gulshan et al., 2016). Furthermore, in freshly isolated mouse bone marrow cells, PIP_2_ is localized on the cell surface (Yoneda et al., 2020). The PIP_2_-binding pocket on the extracellular side of the PAC channel is physiologically unusual. This has been observed once before, in a structure of the TMEM16F channel, which also acts as a lipid scramblase (Feng et al., 2019). Currently, there are no indications that PAC has any lipid scramblase activity.

Considering the recent discovery of PAC and its wide tissue distribution, we speculate that PIP_2_ could be physiologically inhibiting PAC in the outer leaflet of the plasma membrane in specialized cells, or under certain conditions that are currently unknown to us. While the biological function of PAC at the plasma membrane remains a mystery, its primary physiological role is in endosomes and macropinosomes of macrophages. The inner membrane of these intracellular organelles is topologically equivalent to the outer leaflet of the plasma membrane. Some pathogens enter the cells via endocytosis or macropinocytosis and escape degradation in these compartments via fusion of their membrane with the membrane of the organelle. The envelope of some pathogens is enriched in PIP_2_, such as in the human immunodeficiency virus (HIV) (Mücksch et al., 2019). Therefore, we speculate that PIP_2_ could potentially be found in the inner membrane of endosomes during fusion with pathogenic membranes, where it inhibits PAC activity and modulates lumen acidification.

In conclusion, PIP_2_ is the first PAC channel modulator with a characterized binding site and mechanism of action. The advantage of the novel PIP_2_-binding pocket for targeted inhibition of PAC is that it can be exploited for the design of PAC inhibitors that do not have to be cell-permeable. Furthermore, we describe pharmacological properties necessary for PAC inhibition, which include a stable insertion into the membrane and a negative charge that interacts with the positively charged cluster of residues on the pocket. These insights provide a useful tool for the future design of potential therapeutics for acidosis-related diseases implicating the PAC channel.

## Acknowledgements

We thank the Qiu lab for thoughtful discussions. We thank G. Zhao and X. Meng for support with preliminary cryo-EM grid screening at the David Van Andel Advanced Cryo-Electron Microscopy Suite. L.M. is supported by a Boehringer Ingelheim Fonds (BIF) PhD fellowship and National Institute of General Medical Sciences, T32 GM007445 (to the BCMB graduate training program). Z.R. is supported by an American Heart Association (AHA) postdoctoral fellowship (grant 20POST35120556) and the National Institute of Health (NIH) (grant K99NS128258). J. O.-O. is supported by an AHA predoctoral fellowship (grant 18PRE34060025). W.L. is supported by the NIH (grant R01NS112363). Z.Q. is supported by a McKnight Scholar Award, a Klingenstein-Simon Scholar Award, a Sloan Research Fellowship in Neuroscience, and NIH grants (R35GM124824 and R01NS118014). A portion of this research was supported by NIH grant U24GM129547 and performed at the PNCC at OHSU and accessed through EMSL (grid.436923.9), a DOE Office of Science User Facility sponsored by the Office of Biological and Environmental Research.

## Materials and Methods

### Key resources table

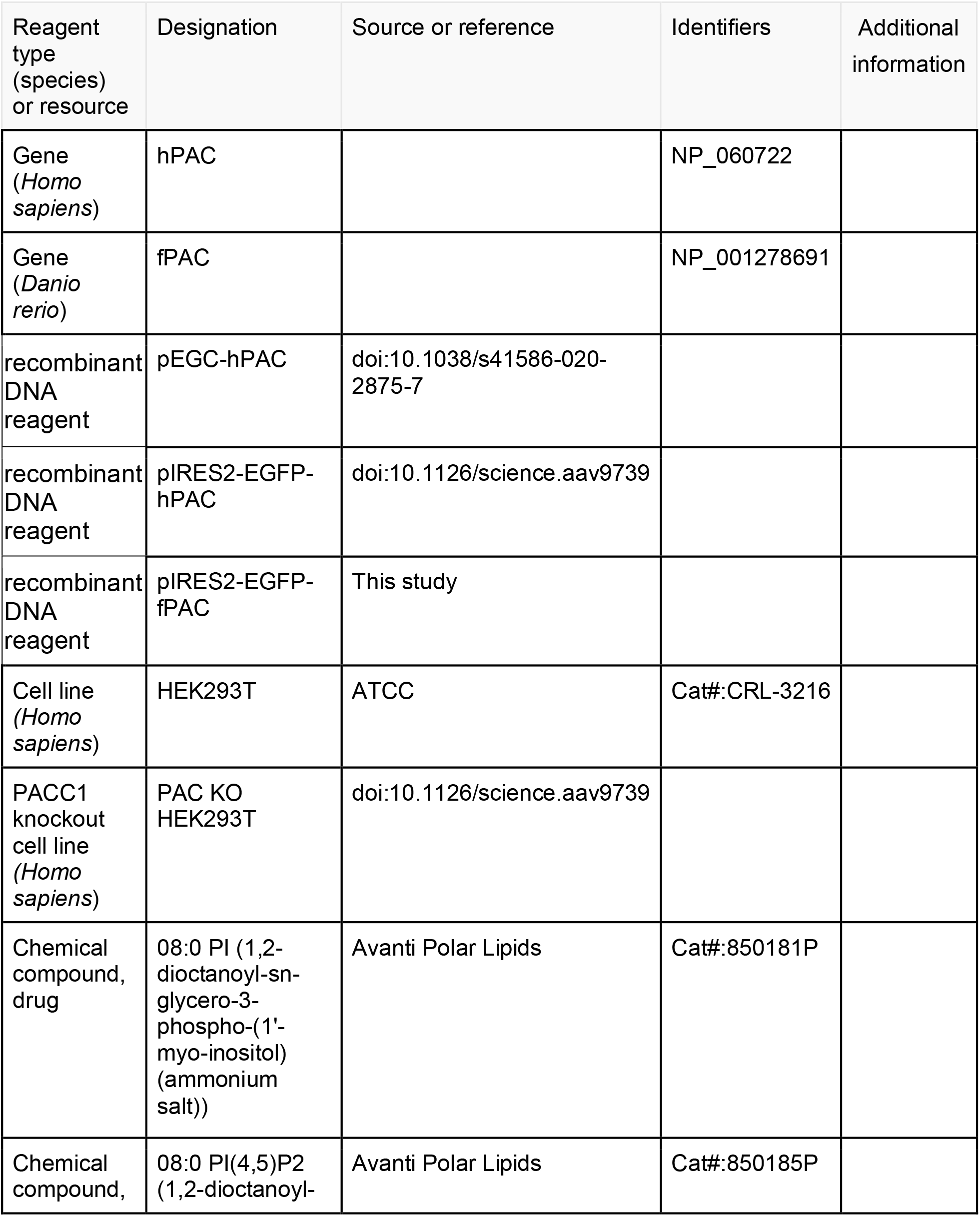
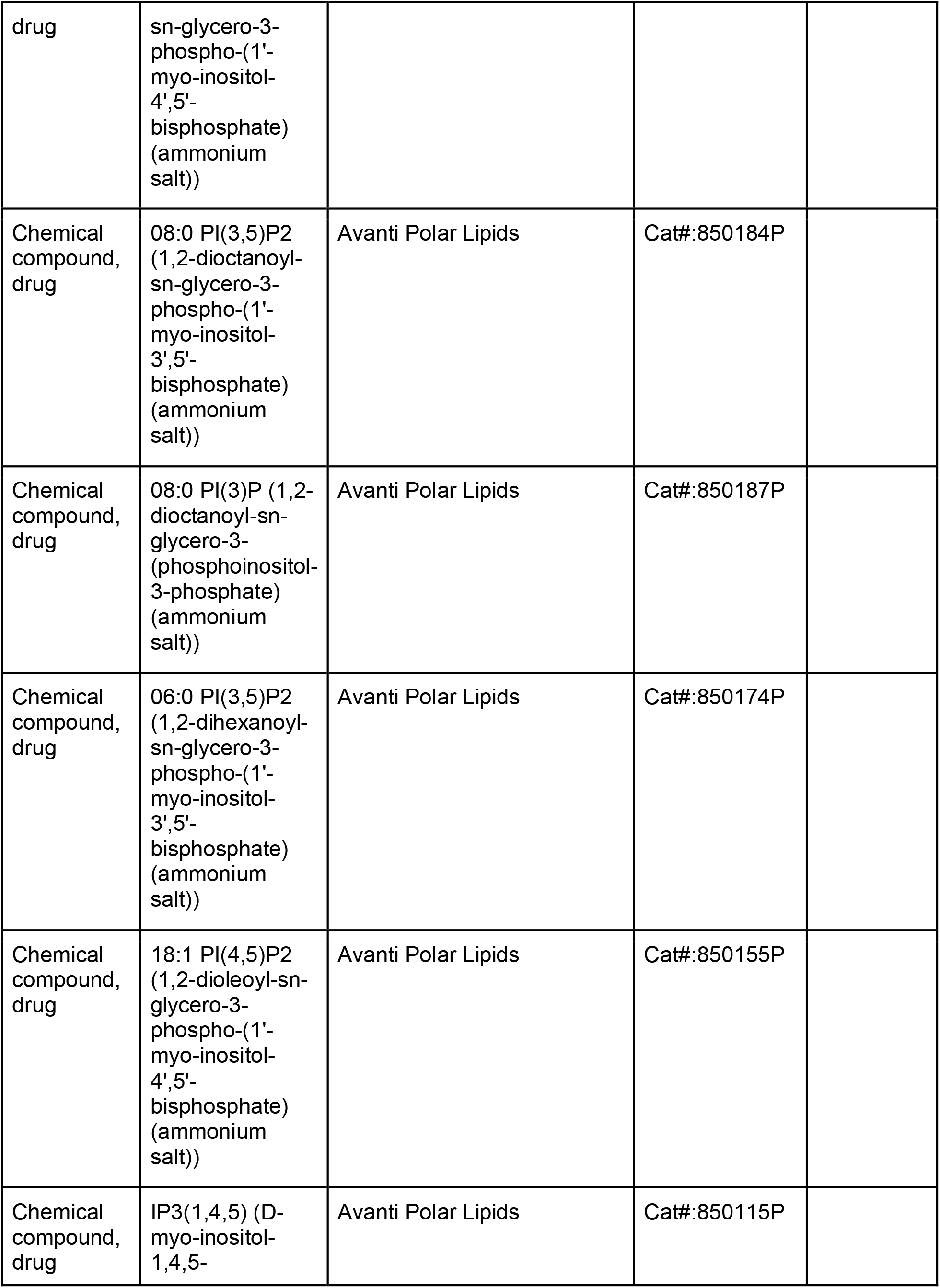
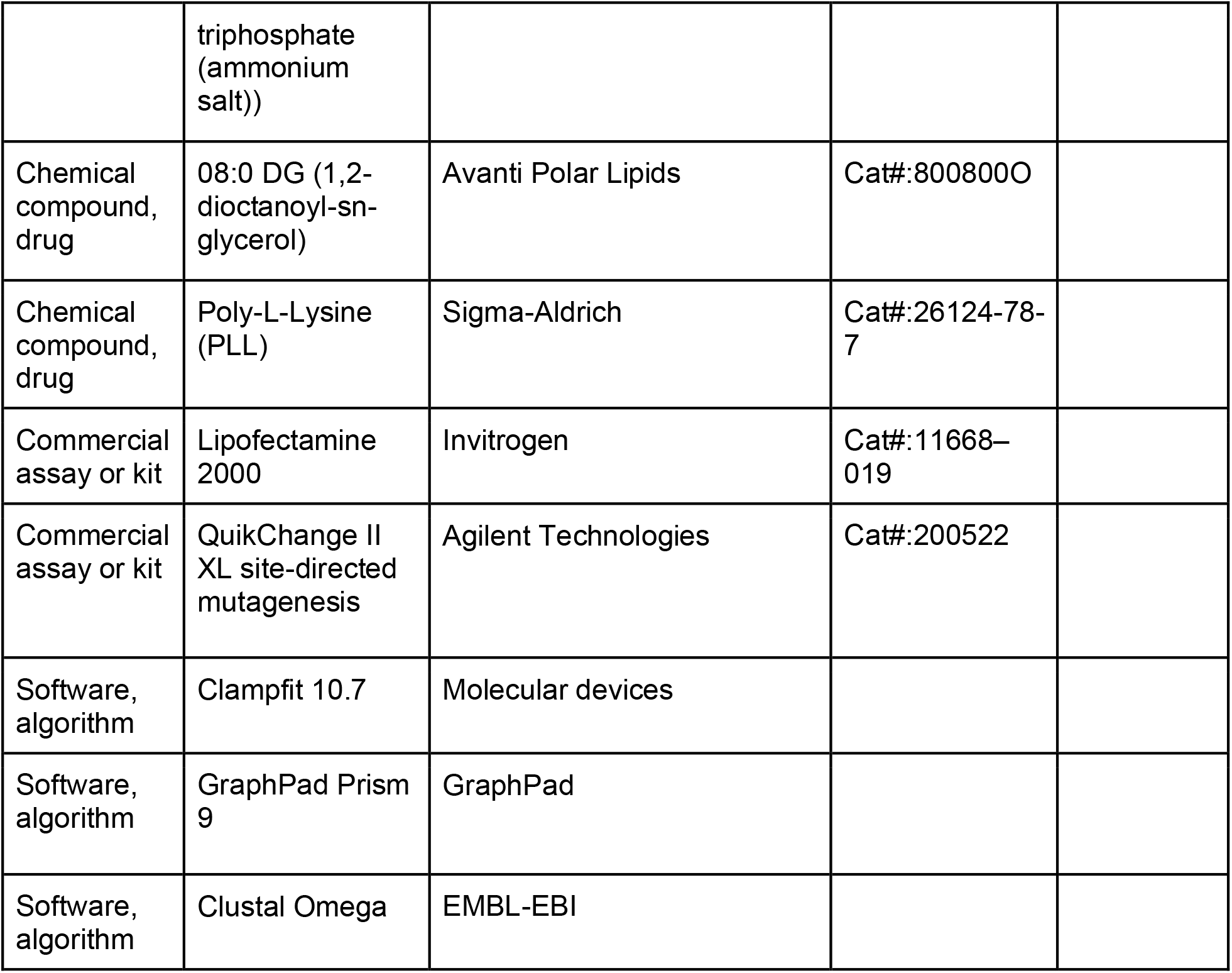

### Cell Culture

HEK293T cells, endogenously expressing PAC channel, or PAC KO HEK293T cells, generated previously using CRISPR technology (Yang et. al, 2019) were maintained in Dulbecco’s modified Eagle’s medium (DMEM) supplemented with 10% fetal bovine serum (FBS) and 1% penicillin/streptomycin (P/S) at 37°C in humidified 95% CO_2_ incubator. PAC KO cells were transfected with 500–800 ng/ml of plasmid DNA using Lipofectamine 2000 (Life Technologies) according to the manufacturer’s instructions. Cells were seeded on 12 mm diameter Poly-l-lysine Sigma-Aldrich) coated glass coverslips and were recorded within 24h after seeding/transfection.

### Constructs and Mutagenesis

Human PAC isoform 2 coding sequence (NP_060722), previously subcloned into pIRES2-EGFP vector (Clontech) using XhoI and EcoRI restriction enzyme sites (Yang et. al, 2019), was used for whole-cell patch-clamp recording experiments. Zebrafish PAC coding sequence (NP_001278691) was subcloned into pIRES2-EGFP vector (Clontech) using NheI and EcoRI restriction enzyme sites. Mutations were introduced using sense and antisense oligos with 15 base pairs of homology on each side of the mutated site. Site-directed mutagenesis was carried out using QuikChange II XL site-directed mutagenesis kit (Agilent Technologies) according to the manufacturer’s instructions. All constructs were confirmed by sequencing the entire open reading frame using Sanger sequencing.

### Sequence Alignments

PAC multiple protein sequence alignments were created using Clustal Omega software (EMBL-EBI). Protein sequences from the following vertebrate species were obtained from UniProt (ID): human PAC (Q9H813), rat PAC (Q66H28), mouse PAC (Q9D771), frog PAC (Q0V9Z3), zebrafish PAC (Q7SY31), bovine PAC (Q2KHV2), orangutan PAC (Q5RDP8), chicken PAC (E1C5B3), and green anole PAC (G1KFB8).

### Lipids and Chemicals

All lipids used in this paper were ordered from Avanti Polar Lipids, and dissolved in water or DMSO, depending on the chain length, to make stock solutions. If not stated otherwise, lipids were added at 10μM concentration directly to the extracellular solution. Please refer to the table for the list of all the lipids used in this paper. Poly-l-Lysine (PLL) (Sigma-Aldrich) was added to the intracellular solution at 100μg/ml.

### Electrophysiology

Whole-cell patch-clamp experiments were performed using the extracellular recording solution (ECS) containing (in mM): 145 NaCl, 2 MgCl_2_, 2 KCl, 1.5 CaCl_2_, 10 HEPES, 10 glucose. The osmolarity of the ECS solution was 300–310 mOsm/kg and the pH was titrated to 7.3 using NaOH. Acidic extracellular solutions contained the same ionic composition, except 5 mM sodium citrate was used as a buffer instead of HEPES, while pH was adjusted using citric acid. ECS solutions were applied 100–200 μm away from the recording cell, using a gravity perfusion system with a small tip. Recording patch pipettes, made of borosilicate glass (Sutter Instruments), were pulled with a Model P-1000 multi-step puller (Sutter Instruments). The patch pipettes had a resistance of 2–4 MΩ when filled with an intracellular solution (ICS) containing (in mM): 135 CsCl, 2 CaCl_2_, 1 MgCl_2_, 5 EGTA, 4 MgATP, 10 HEPES. The osmolarity of the ICS solution was 280–290 mOsm/kg and pH was titrated to 7.2 using CsOH. *I*_Cl,H_ recordings were acquired using voltage ramp pulses from –100 to +100 mV. The time interval between two ramp pulses was 2 or 5 s at a speed of 1 mV/ms and the holding potential was 0 mV. All recordings were performed with a MultiClamp 700B amplifier and 1550B digitizer (Molecular Devices) at room temperature. Signals were filtered at 2 kHz, digitized at 10 kHz, and the series resistance was compensated for at least 80% (Yang et al., 2019).

### Data Analysis

Electrophysiology data were analyzed using Clampfit 10.7. Statistical analysis was performed using GraphPad Prism 9 software. Comparison between two groups was carried out using an unpaired two-tailed Student’s *t* test unless stated otherwise. Multiple group comparisons were performed using ordinary one-way analysis of variance (ANOVA). The significance level was set at p < 0.05. All numerical data are shown as mean ± SEM. For the time-constant experiments, the currents were fit using a one-phase decay equation: Y=(Y0 - Plateau)*exp(-K*X) + Plateau, where Y0 was the time-point of adding PIP_2_ to the cells. For the IC_50_ values, the normalized data was fitted to the following sigmoidal 4PL equation, where X is log (concentration): Span = Top - Bottom; Y=Bottom + (Top-Bottom)/(1+10^((LogIC_50_-X)*HillSlope)).

### Protein Expression and Purification

The pEGC-hPAC plasmid containing the human PAC gene, a Strep-tag II tag, a thrombin cleavage site, an eGFP, and an 8xHis tag, was used for expressing PAC protein in mammalian cells using baculovirus (Goehring et al., 2014). The bacmid was produced by transforming the DH10Bac cells and positive white clones were selected from a Luria Broth (LB) plate with kanamycin (50 μg/mL), tetracycline (10 μg/mL), and gentamicin (7 μg/mL) resistance. Bluo-gal (100 μg/mL Bluo-gal) and IPTG (40 μg/mL) were also included in the plate for the selection of recombinant bacterial clones. Bacmid DNA was purified from LB cultures of the white colonies using the alkaline lysis method. The bacmid was then transfected into adherent Sf9 cells grown in Sf-900 II media (Gibco) using Cellfectin II reagent by using the manufacturer’s recommended protocols. After 5 days, the media of Sf9 cell culture was filtered as the P1 virus. Subsequently, the P2 virus was made by infecting suspension Sf9 cells grown in Sf-900 II media with P1 virus at a 1:5000 ratio (v/v). After another 5 days, the P2 virus was harvested, filtered, and stored at 4°C with 1% fetal bovine serum (FBS). Mammalian cells (tsA-201 cell line) grown in FreeStyle 293 media (Gibco) supplemented with 1% FBS were used for protein expression. When cells reached 3.5×10^6 cells/ml density, 10% (v/v) P2 virus was added to tsA-201 cells and cells were allowed to grow for another 8-12 h at 37°C. To boost protein expression, 5 mM sodium butyrate was added to the cell culture and cells allowed to grow for 60 h at 30°C. The infected cells expressing PAC were then spun down and the pellet stored at −80°C until protein purification.

The cell pellet was resuspended in ice-cold TBS buffer (20mM Tris pH8 and 150mM NaCl) with a protease inhibitor cocktail (1 mM PMSF, 0.8 μM aprotinin, 2 μg/ml leupeptin, 2 mM pepstatin A) and lysed by sonication. The debris was removed by centrifugation at 4000 rpm for 10 min at 4°C. The supernatant underwent ultracentrifugation at 40,000 rpm for 1 h and the cell membranes were collected. The membranes were solubilized in a TBS buffer with 1% glyco-diosgenin (GDN) detergent (Anatrace) and the protease inhibitor cocktail for 1 h at 4°C with gentle rotation. The sample was clarified by ultracentrifugation at 40,000 rpm for 1 h. The supernatant was subjected to immobilized metal affinity chromatography (IMAC) with talon resin (Takara Bio USA). The bound protein was washed with TBS buffer containing 0.02% GDN and 20 mM imidazole and eluted with TBS buffer containing 0.02% GDN and 250 mM imidazole. The PAC protein was then concentrated to 1 ml using a 100 kDa concentrator. The sample was then mixed with soybean lipid extract (Anatrace) and His-tag free membrane scaffold protein 1E3D1 (Denisov et al., 2007) at a 1:200:3 molar ratio. The GDN detergent was removed through three rounds of biobeads (Bio-Rad) incubation at 4°C. To remove “empty” nanodiscs, the sample was filtered to remove biobeads and incubated with talon resin at 4°C for another 1 h. The volume of the sample was expanded to 25 ml by adding a TBS buffer such that the imidazole concentration was lowered to 10 mM. The resin was washed with a TBS buffer containing 10 mM imidazole, and the protein was eluted by a TBS buffer with 250 mM imidazole. PAC-nanodisc protein was then concentrated to 500 μL using an Amicon Ultra-15 concentrator (100 kDa cutoff). Thrombin (0.03 mg/ml) was added to cleave GFP from the PAC protein at 4°C overnight. PAC-nanodisc was further purified by size-exclusion chromatography (SEC) using TBS buffer. The peak fractions were concentrated to 5 mg/ml prior to making cryo-EM grids.

### Cryo-EM Grid Preparation

Purified human PAC protein in nanodiscs were first mixed with 1 mM diC_8_-PI(4,5)P_2_ (Avanti) on ice for 1 h. The pH of the protein sample was adjusted to 4.0 by adding acidic acid buffer (1M, pH 3.5) at 1:20 ratio (v/v). We also added 0.5 mM fluorinated octyl maltoside (Anatrace) to improve sample quality. An FEI Vitrobot Mark III was used for plunge-freezing. Specifically, a 3-μl aliquot of the protein sample was applied to a glow-discharged Quantifoil holey carbon grid (Au 300 2/1 mesh) (Electron Microscopy Sciences), blotted for 2 s, vitrified in liquid ethane, and transferred to liquid nitrogen for storage. The temperature and humidity of the chamber was kept at 18 °C and 100% throughout the grid preparation.

### Cryo-EM Data Collection

The cryo-EM grids were initially screened in an FEI Talos Arctica transmission electron microscope equipped with a K2 summit camera. High-resolution data collection was facilitated by the Pacific Northwest Center for Cryo-EM (PNCC) using an FEI Titan Krios transmission electron microscope equipped with a BioQuantum energy filter (20 eV slit width) and a K3 camera with a nominal magnification of 105,000. SerialEM was used for automated data collection in super-resolution mode with a pixel size of 0.413Å (Mastronarde, 2005). The raw movie stack contained a total of 52 frames with a total dose of 50 e^−^/Å^2^. The nominal defocus value was allowed to vary between −0.6 to −2.4 μ m.

### Cryo-EM Data Processing

The cryo-EM data processing workflow is summarized in Figure S3. Specifically, the raw movies were motion corrected using relion 3.1 and binned to the physical pixel size at 0.826Å (Zivanov et al., 2018). The defocus parameters of motion-corrected micrographs were estimated using ctffind 4.1.10 (Rohou & Grigorieff, 2015). Particle picking was performed using both gautomatch_v0.56 (https://www2.mrc-lmb.cam.ac.uk/research/locally-developed-software/zhang-software/) and topaz v0.2.5 (Bepler et al., 2019). Particles picked by each program were independently subjected to 2D classification (relion 3.1) or heterogeneous refinement (cryosparc v3.0) to get rid of junk particles (Zivanov et al., 2018, Punjani et al., 2017). Good particles with clear features were pooled together and refined with C3 symmetry in relion 3.1. 3D refinement with a solvent mask, resulting in a 4.4 Å map. We noticed that the size of nanodiscs could be heterogeneous, which may negatively affect particle alignment. Therefore, we created a loose mask of the protein based on the atomic model and performed signal subtraction to remove the nanodisc signal. The process allowed us to obtain a reconstruction at 4.2 Å resolution. To sort out conformational heterogeneity of the dataset, we performed 3D classification without image alignment in relion 3.1. The best class of the job was selected and refined to 3.6 Å resolution. We then performed several rounds of CTF refinement and Bayesian polishing (Zivanov et al., 2019), and the map was eventually refined to 3.17 Å. We noticed an improvement in the map quality when the box size of the images was expanded from 240 pixels to 300 pixels at this stage. To further improve map reconstruction, we first split the original consensus particles after 2D and heterogeneous refinement into 6 portions. We combined each portion with the best particles that gave rise to the 3.17 Å reconstruction and performed another round of 3D classification. This procedure was effective in attracting good particles from the initial consensus map particles. After combining the best class and removing duplicates, we identified 84k particles that could be refined to 3.07 Å in relion after iterative CTF refinement and Bayesian Polishing. We then exported the particles to cryosparc and conducted CTF refinement followed by local refinement. We also supplied a mask to get rid of nanodisc signal during the refinement. In the end, we obtained a 2.71 Å reconstruction as judged by gold standard Fourier shell correlation.

### Model Building, Validation, and Analysis

The atomic model was generated by first docking the structural model of human PAC at pH 4.5 (PDBID: 7SQH) into the cryo-EM map (Wang et al., 2022). The diC_8_-PI(4,5)P_2_ molecule was manually placed into the cryo-EM density. The Grade Web Server (Bepler et al., 2019) was used to generate a restraint file for flexible fitting of diC_8_-PI(4,5)P_2_ molecule. Subsequently, the model underwent real space refinement in phenix and manual adjustment to fix Ramachandran outliers, rotamer outliers, and clashes (Adams et al., 2010). The final model was validated by the molprobity in phenix to obtain validation statistics (Williams et al., 2018). The cryo-EM map and atomic model were visualized using UCSF ChimeraX (Pettersen et al., 2021). The pore profile of PAC channel was calculated using HOLE 2.0 program (Smart et al., 1996).

## Supplementary figures

**Figure S1.**
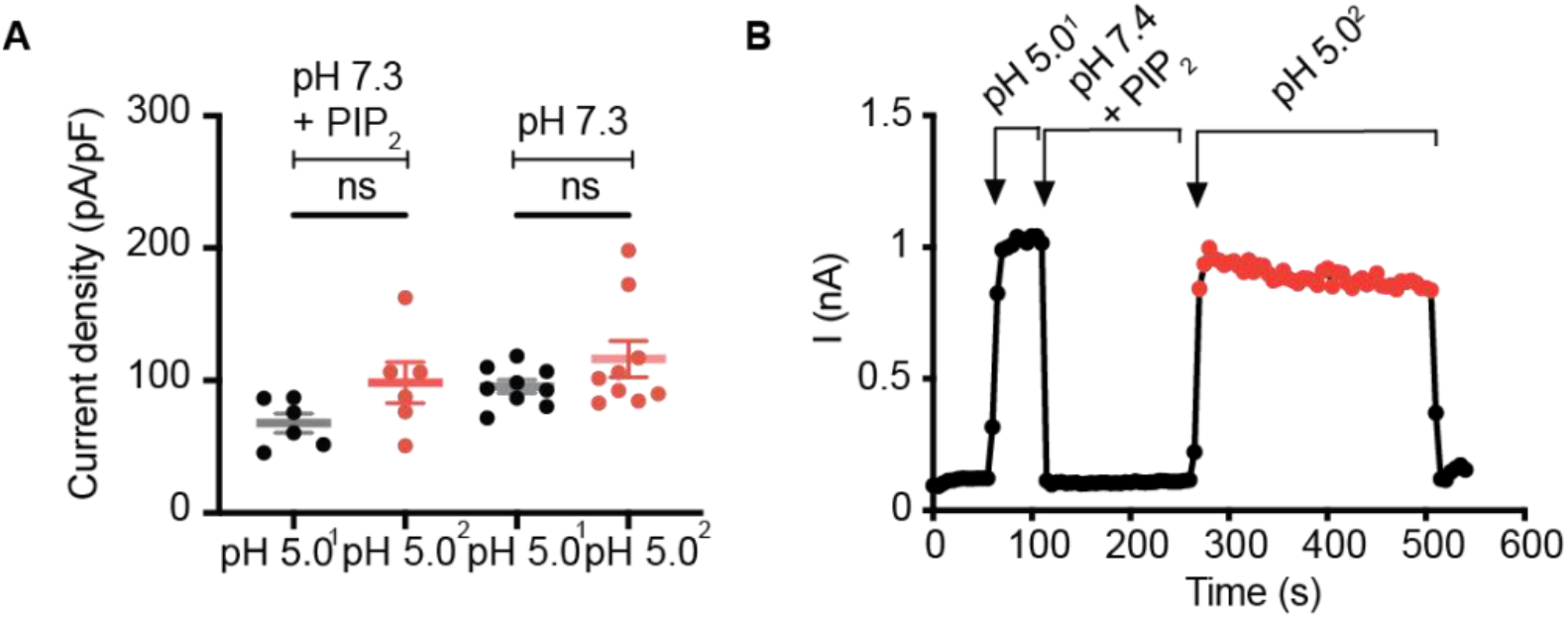
PIP_2_ doesn’t bind to the closed PAC channel. **(A)** Current amplitude was measured at pH 5.0 before (1) and after (2) perfusion of diC_8_-PIP_2_ at pH 7.3 (+PIP_2_). The control cells were perfused with pH 7.3 only (-PIP_2_). There was no significant difference in current density before and after treatment with PIP_2_ at neutral pH. Statistical significance was determined using a two-tailed Student’s unpaired *t* test. ns: not significant. **(B)** Representative current trace (3s/sweep) of the experiment described in A).

**Figure S2.**
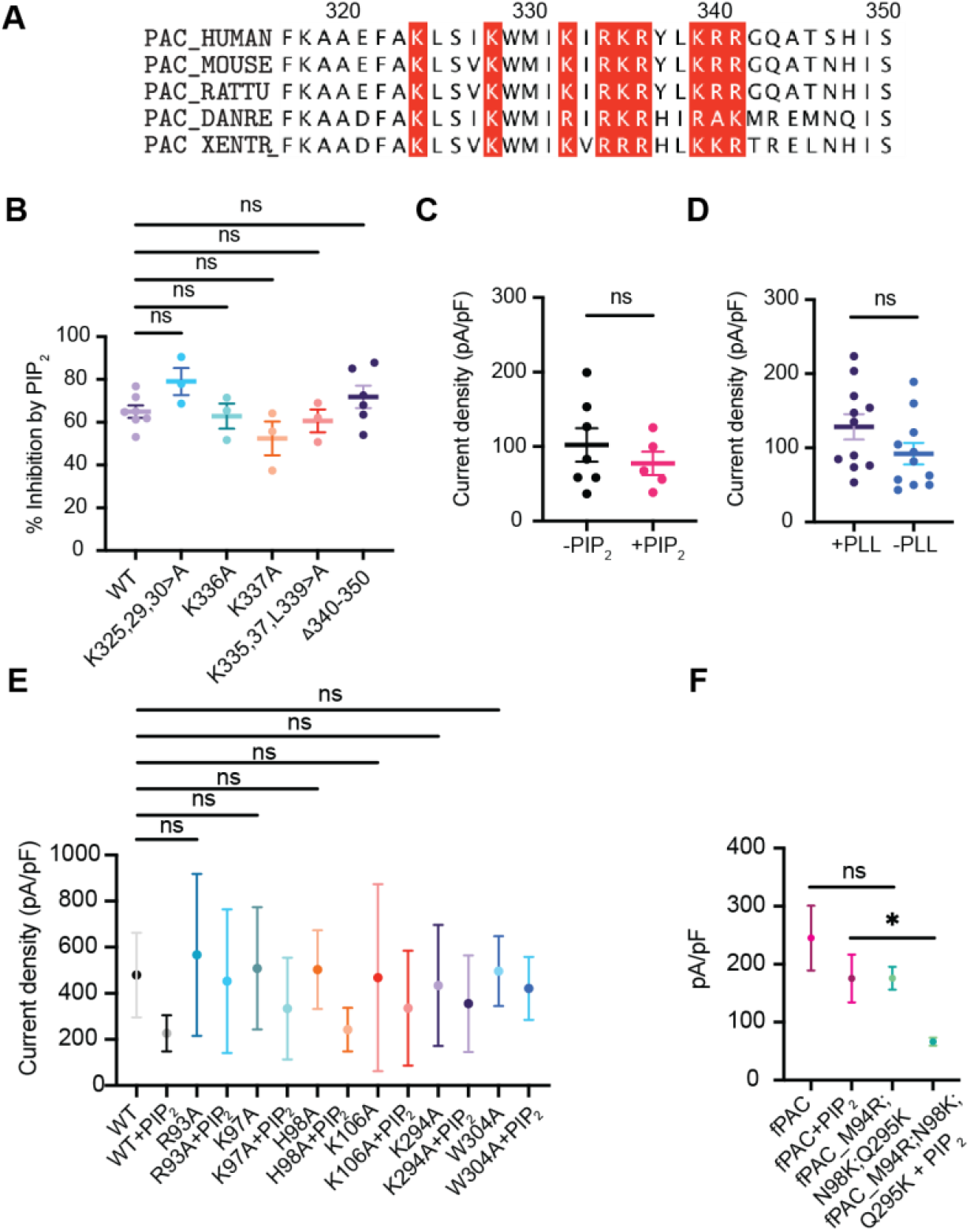
The binding site for PIP_2_ is not on the intracellular side of the PAC channel. **(A)** We hypothesized that a cluster of positively charged residues at the C-terminus of PAC, outlined in red, binds PIP_2_. **(B)** Potential cytosolic PIP_2_-binding residues were screened by making grouped alanine mutations, or single alanine mutants, and by deleting a 10 amino-acid sequence at the C-terminal end of PAC, but there was no significant difference in PAC inhibition by PIP_2_. Statistical significance was determined using one-way ANOVA with Dunnett post hoc test; ns: not significant. Bars are reported as mean ± SEM. **(C)** 10μM diC_8_-PIP_2_ added through a patch pipette containing intracellular solution (ICS) in whole-cell configuration showed no significant difference when compared to ICS without PIP_2_. Statistical significance was determined using a two-tailed Student’s unpaired t test. ns: not significant. Bars are reported as mean ± SEM. **(D)** There was no significant difference before and after depleting endogenous PIP_2_ from the inner leaflet by applying 100μg/ml of Poly-l-Lysine (PLL) through the patch pipette. Statistical significance was determined using a two-tailed Student’s unpaired t test. ns: not significant. Bars are reported as mean ± SEM. **(E)** Current density before adding PIP_2_ is unaffected when PIP_2_-binding residues are mutated to alanine in hPAC. Statistical significance was determined using one-way ANOVA with Dunnett post hoc test. ns: not significant. Bars are reported as mean ± SEM. **(F)** Current density is unchanged when fPAC residues are mutated to the corresponding hPAC residues, while it is significantly decreased in the presence of PIP_2_. Statistical significance was determined using a two-tailed Student’s unpaired t test. ns: not significant. **p < 0.01. Bars are reported as mean ± SEM.

**Figure S3.**
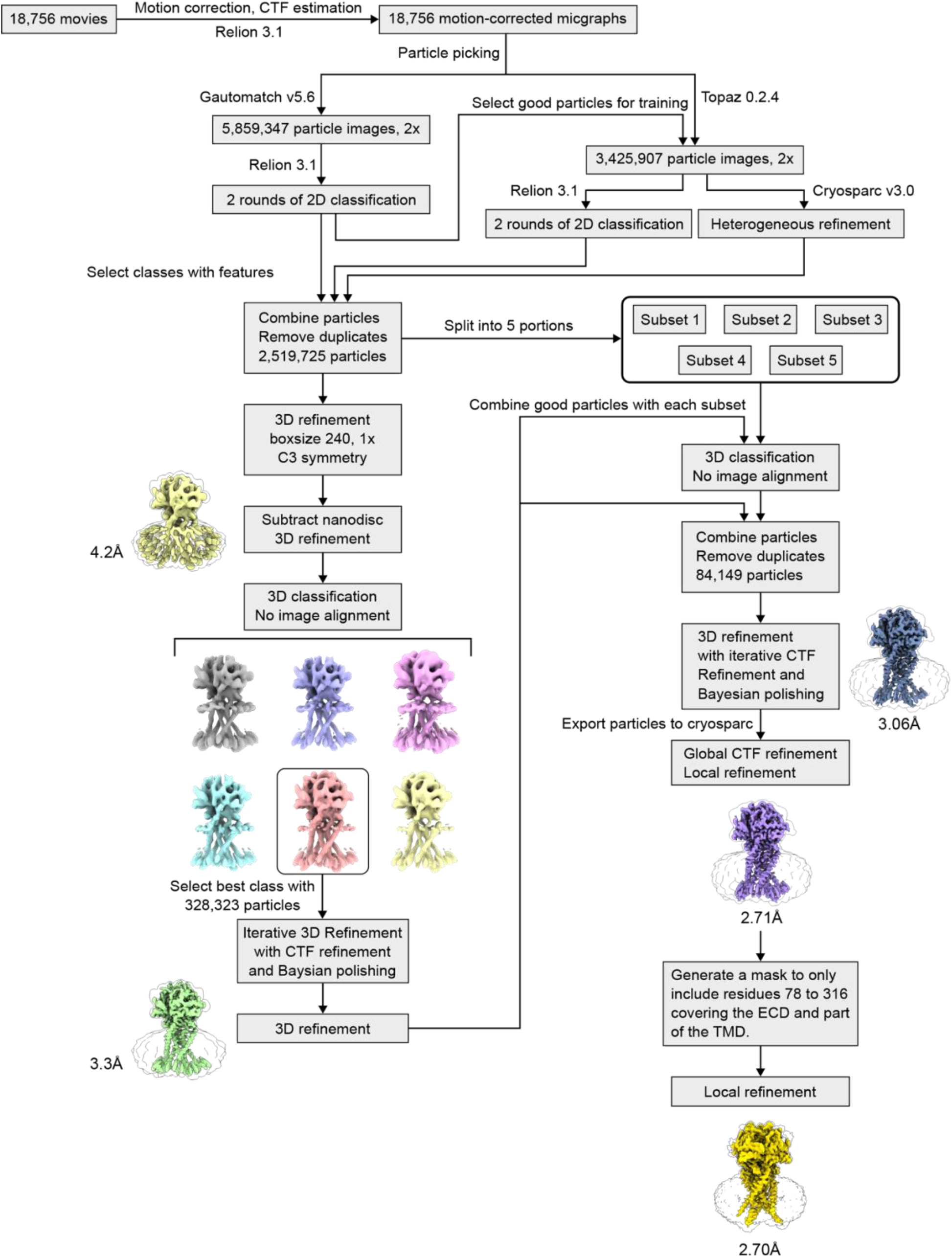
The cryo-EM data processing workflow of human PAC in nanodisc at pH 4.0 with 0.5 mM PIP_2_ dataset. A more detailed description of this process can be found in the method section.

**Figure S4.**
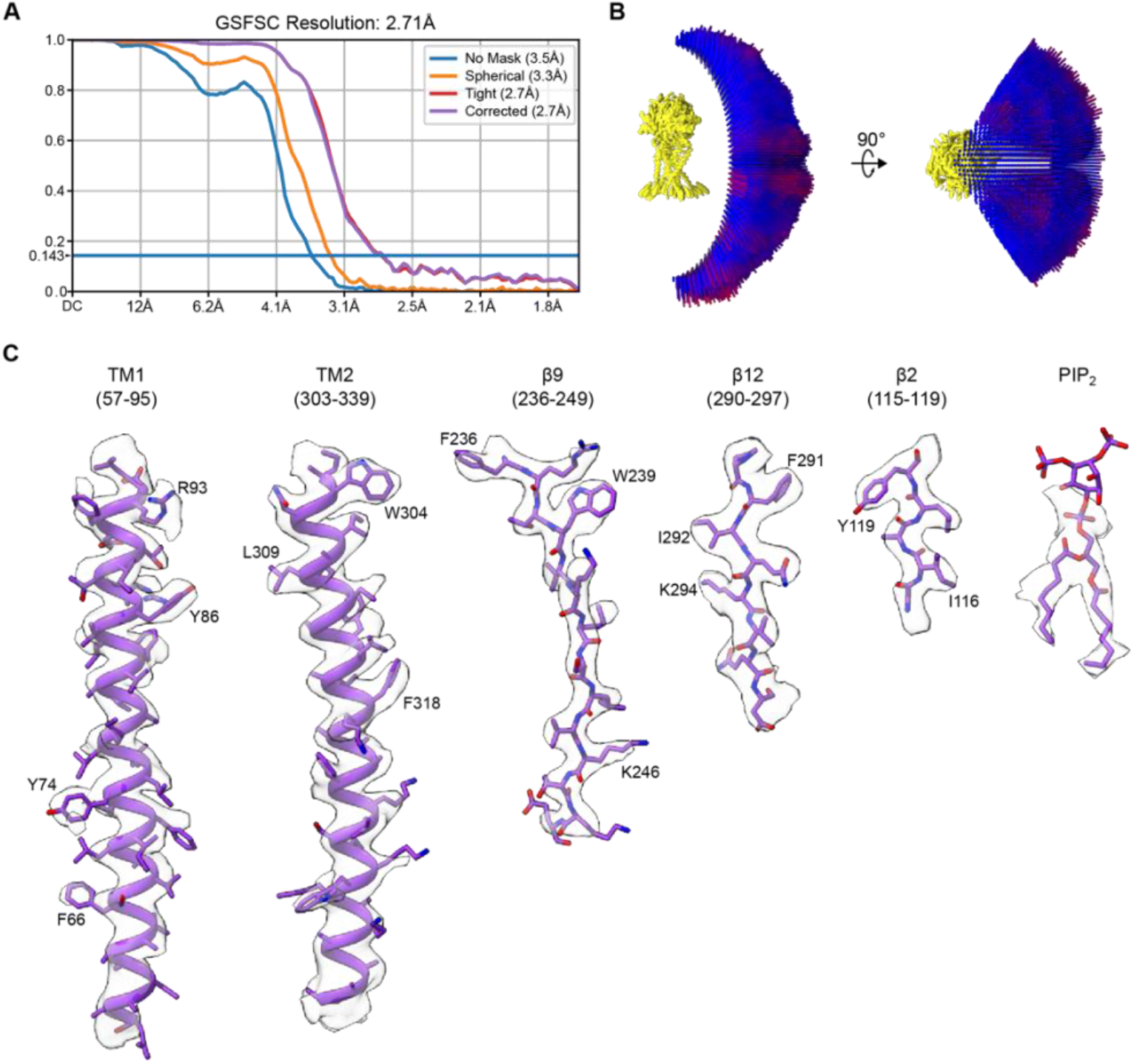
The reconstruction metrics of human PAC in nanodisc at pH 4.0 with 0.5 mM PIP_2_. **(A)** The gold-standard Fourier shell correlation curve of the final cryo-EM map. **(B)** The angular distribution of particles that give rise to the final reconstruction. **(C)** The representative densities of the reconstruction map, including TM1/2, β2, β9, β12, and PIP_2_.

**Table S1:** Cryo-EM data collection, refinement, and validation statistics

| Human PAC with PI(4,5)P<sub>2</sub> at pH 4.0 |  |
| --- | --- |
| **Data collection and processing** |  |
| Magnification | 105,000 |
| Voltage (kV) | 300 |
| Electron exposure (e<sup>-</sup>/Å<sup>2</sup>) | 50 |
| Defocus range (μm) | −0.6 to −2.4 |
| Pixel size (Å) | 0.826 |
| Symmetry imposed | C3 |
| Initial particle images (no.) | 5,859,347 |
| Final particle images (no.) | 84,149 |
| Map resolution (Å) | 2.71 |
| FSC threshold | 0.143 |
| Map resolution range (Å) | 2.71 – 246.2 |
| **Refinement** |  |
| Initial model used (PDB code) | 7SQH |
| Model resolution (Å) | 3.27 |
| FSC threshold | 0.5 |
| Map sharpening *B* factor (Å<sup>2</sup>) | −95 |
| Model composition |  |
| Non-hydrogen atoms | 6930 |
| Protein residues | 849 |
| Ligands | 15 |
| R.m.s. deviations |  |
| Bond lengths (Å) | 0.011 |
| Bond angles (°) | 1.621 |
| Validation |  |
| MolProbity score | 1.88 |
| Clashscore | 7.26 |
| Poor rotamers (%) | 2.34 |
| Ramachandran plot |  |
| Favored (%) | 96.80 |
| Allowed (%) | 3.20 |
| Disallowed (%) | 0.00 |

